# Orthogonal neural codes for phonetic features in the infant brain

**DOI:** 10.1101/2021.03.28.437156

**Authors:** Giulia Gennari, Sébastien Marti, Marie Palu, Ana Fló, Ghislaine Dehaene-Lambertz

## Abstract

Creating invariant representations from an ever-changing speech signal is a major challenge for the human brain. Such an ability is particularly crucial for preverbal infants who must discover the phonological, lexical and syntactic regularities of an extremely inconsistent signal in order to acquire language. Within visual perception, an efficient neural solution to overcome signal variability consists in factorizing the input into orthogonal and relevant low-dimensional components. In this study we asked whether a similar neural strategy grounded on phonetic features is recruited in speech perception.

Using a 256-channel electroencephalographic system, we recorded the neural responses of 3-month-old infants to 120 natural consonant-vowel syllables with varying acoustic and phonetic profiles. To characterize the specificity and granularity of the elicited representations, we employed a hierarchical generalization approach based on multivariate pattern analyses. We identified two stages of processing. At first, the features of manner and place of articulation were decodable as stable and independent dimensions of neural responsivity. Subsequently, phonetic features were integrated into phoneme-identity (i.e. consonant) neural codes. The latter remained distinct from the representation of the vowel, accounting for the different weights attributed to consonants and vowels in lexical and syntactic computations.

This study reveals that, despite the paucity of articulatory motor plans and productive skills, the preverbal brain is already equipped with a structured phonetic space which provides a combinatorial code for speech analysis. The early availability of a stable and orthogonal neural code for phonetic features might account for the rapid pace of language acquisition during the first year.

**SIGNIFICANCE STATEMENT:** For adults to comprehend spoken language, and for infants to acquire their native tongue, it is fundamental to perceive speech as a sequence of stable and invariant segments despite its extreme acoustic variability. We show that the brain can achieve such a critical task thanks to a factorized representational system which breaks down the speech input into minimal and orthogonal components: the phonetic features. These elementary representations are robust to signal variability and are flexibly recombined into phoneme-identity percepts in a secondary processing phase. In contradiction with previous accounts questioning the availability of authentic phonetic representations in early infancy, we show that this neural strategy is implemented from the very first stages of language development.

## INTRODUCTION

A major, fundamental challenge for any brain is to build stable representations of a changing world. In particular regarding speech, the subtle phonetic distinctions between “bog”, “dog” or “big” must be perceived steadily despite the large acoustic differences separating the raspy voice of a whispering elderly man and the fluty screams of a little girl. Since the richness of the human lexicon is based on fine phonetic differences of this sort, how infants come to discover them in a highly variable signal has long been the subject of debate.

Recent proposals, based on neuronal recordings during object (Behrens et al., 2018) and face recognition (L. Chang & Tsao, 2017), suggest that in order to deal with signal inconsistency, the brain factorizes the input into independent and orthogonal low-dimensional components, each coding for a different dimension of variation. The components are thought to be subsequently recombined to yield unified percepts. Can such an account be applied to speech? Apart from any neural consideration, linguists have proposed an abstract definition of phonemes as bundles of orthogonal elementary features, each corresponding to a binary code that summarizes an articulatory dimension and its acoustic correlates (Halle, 2013). For instance, the phonemes “b” and “d” from the example above share all parameters (+consonantal and -vocalic, +obstruent and -sonorant, +voiced, etc.) except for the place of articulation (+labial/-alveolar vs. +alveolar/-labial). Given their linguistic characteristics (distinctive, minimal and combinable), these features might correspond to the basic decomposition axes harnessed by the brain to overcome speech variability. In the last years, high-resolution intracranial recordings on adults (Mesgarani et al., 2014) and fMRI adult data (Arsenault & Buchsbaum, 2015) have provided evidence in line with this hypothesis: a partial neural specialization for phonetic features was observed during passive listening of speech.

For what concerns infants however, vocal production develops slowly during the first year through vocal plays in an effort to imitate ambient language (Kuhl & Meltzoff, 1996). Babbling, which signals the beginning of a relatively controlled articulation, enriches vocalizations only from the second semester. Given their initial inability to produce most phonemes, can preverbal infants use a code originally defined by articulatory gestures? The prevailing view rejects such an eventuality. During the first semester infants are thought to analyze speech merely along domain-general spectrotemporal dimensions (Kuhl, 2004). They would gradually converge to an adult-like phonetic space only later, through native language exposure and the motor feedback provided by the progressive acquisition of articulatory/motor skills (Kuhl et al., 2008; Westermann & Reck Miranda, 2004; Vilain et al., 2019). Yet, from birth on, infants are capable of perceiving stable speech segments despite acoustical variations. For instance, they identify phonemes independently of the speaker (Dehaene-Lambertz & Pena, 2001) or the co-articulation context (Mersad & Dehaene-Lambertz, 2016) and can track phonologic information across changes in prosody (Fló et al., 2019). Moreover, words start to be stored at 6 months, thus earlier than predicted by mainstream accounts (Tincoff & Jusczyk, 1999; Bergelson & Swingley, 2012). By revealing the capacity to overcome signal variability, these observations prompt to reconsider the format of early speech encoding. We hypothesized that pre-babbling infants factorize the speech signal along independent low-dimensional components, creating a structured phonetic space. The latter would *(a)* account for the refined linguistic abilities documented in very young infants (Dehaene-Lambertz & Gliga, 2004); and *(b)* facilitate the discovery of phonetic regularities beyond surface differences thereby providing the ideal basis for lexicon acquisition.

To test our proposal, we exposed twenty-five 3-month-old infants to 120 natural consonant-vowel syllables and examined their event-related brain potentials (ERPs) using time-resolved multivariate pattern analyses. We first assessed whether linear classifiers could separate neural responses according to phonetic distinctions. We then examined how decoders trained on particular data subsets performed once a given variation was controlled. this analytical procedure was crucial to the aim of the study, in that the level of generalization beyond the training set enabled to determine the precise format of the neural codes underlying decodability (Kriegeskorte & Douglas, 2019) At a minimum, we expected linguistic and speaker information to be encoded in parallel, as suggested by previous behavioral and EEG studies (Kuhl, 1979; Dehaene-Lambertz & Pena, 2001). In other words, we expected estimators trained on ERPs to syllables produced by a female voice to obtain similar results when tested on ERPs to syllables pronounced by a male speaker. We then examined generalization performances across co-articulatory contexts (e.g. are syllables containing “i” and “o” processed through a common consonantal code?) and across featural dimensions (e.g. are both the obstruent “b” and the sonorant “m” encoded as “labial”?). Only a complete generalization along all these steps can assure that infant speech encoding is ultimately based on phonetic features. Furthermore, tracing the time course of the generalization patterns and analyzing class confusability gave us the opportunity to elucidate whether consonants and syllables were deconstructed into, or reconstructed from, elementary parts.

## RESULTS

Experimental sessions lasted about 1 hour with a total of ~3100 stimuli presented to each baby. Syllables were chosen to independently vary the consonantal dimensions of manner (obstruent vs. sonorant) and place of articulation (labial vs. alveolar vs. velar). Each consonant was coupled with two vowels (/i/ and /o/) and produced by a male and a female speaker in five distinct utterances to ensure acoustic and co-articulatory variability across tokens with the same phonetic profile (Figure 1A).

**Figure 1:**
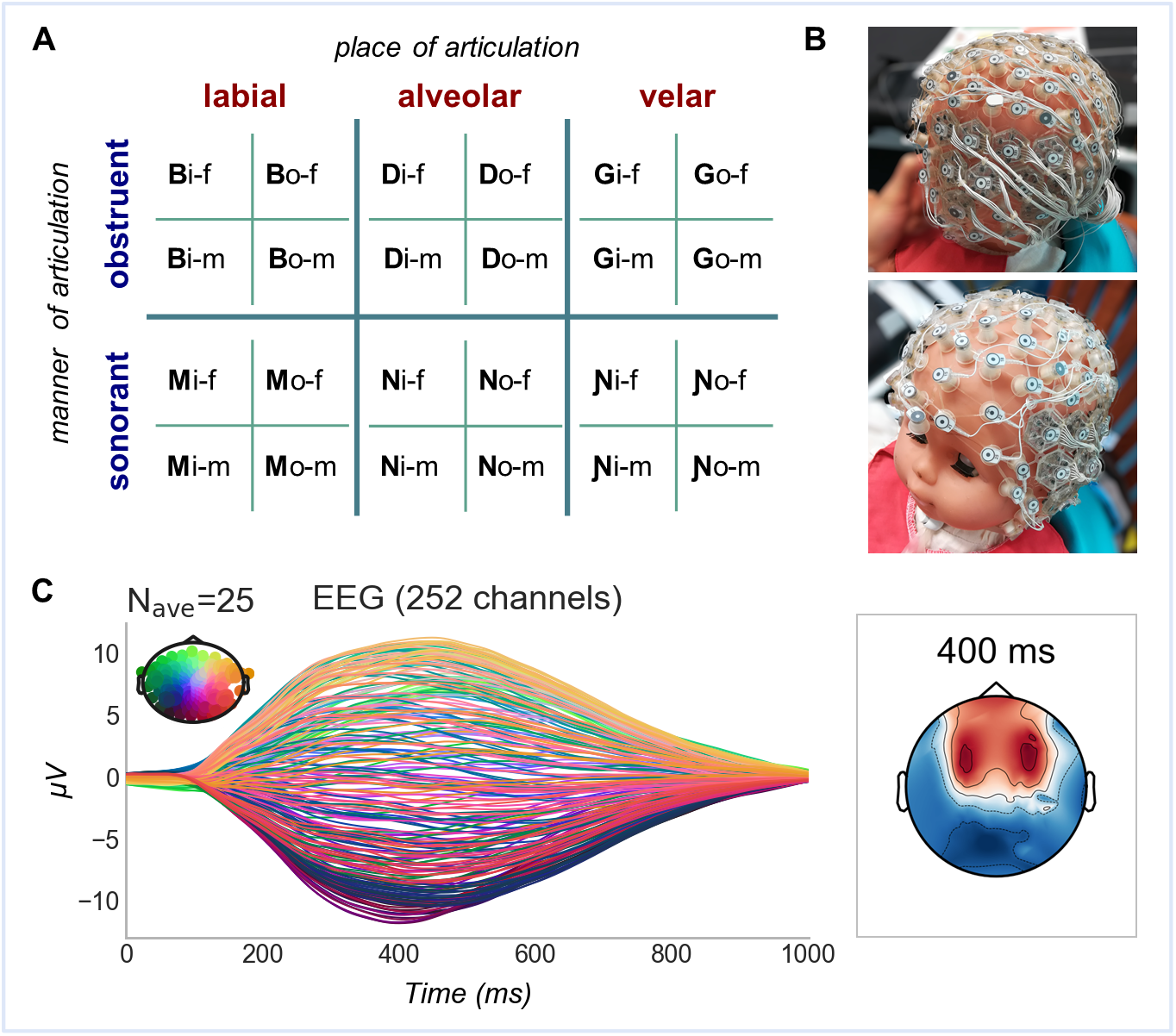
Experimental set-up and average syllable-related potential. (A) Stimuli sub-conditions and their phonetic characteristics (f=female, m=male voice). (B) 256 channels super-high-density net on the head of an infant manikin: tight grids of custom electrodes are arranged over the auditory linguistic areas of the superior temporal lobe (see also Figure S2). (C) Grand average ERP: all conditions are pulled together.

The dimensions of manner and place of articulation were chosen due to the different levels of consistency characterizing their acoustic correlates: whereas manners are reflected in prominent spectrotemporal prototypes (Stevens, 2000), the acoustic cues for place are more subtle (Shannon et al., 1995) and complex (Smits et al., 1996), hence fundamentally dependent on the context of production (Fowler, 1994). Following these observations, our ability to detect stable place contrasts across different production circumstances is commonly seen as the ultimate challenge to address in order to understand human speech perception. Intriguingly, although able to form place-based categories, animals have been shown to process place contrasts in a context-dependent way (Sinnott & Gilmore, 2004). An initial investigation of the similarity structure embedded in our stimuli set (Figure S1) confirmed the diverging nature of manner and place acoustic cues. Along an average sound duration of 400ms, the auditory pairwise dissimilarity of the stimuli was best described by manner of articulation distinctions up to 140ms (i.e. during the consonantal portion) and later by the vowel (Figure S1D). Acoustic similarities were additionally shaped by voice gender throughout the entire syllable, while they did not have any straightforward relationship with the place of articulation (Figure S1D).

Infant ERPs were recorded with a high-density custom net featuring 256 channels (Figures 1B and S2; see also Figure 1C for the grand average across all syllables). While the intensive electrode coverage combined with the thinness of infant skulls maximized the spatial resolution of our recordings, univariate analyses are poorly suited for separating the activity of neuronal clusters that are spatially close. We thus opted for a more powerful multivariate analysis approach (Stokes et al., 2015): we trained and tested series of linear estimators on brief (20ms) consecutive windows all along the high-density ERPs. Our goal was to define the granularity of the infant coding scheme for speech: is it syllabic, phonetic or featural?

### Successful classification is achieved on the basis of dynamic and discrete neural patterns

We first assessed whether infant neural responses were separable according to phonetic classes. Figure 2A-B show that obstruents were distinguished from sonorants starting from 80ms after syllable onset (pclust =0.0001; peak performance observed at 200ms: AUC=0.735±0.08, chance= 0.5), while places of articulation were reliably classified over two time windows: 220-480ms (pclust=0.0001; peak at 260ms: M=0.545±0.039); and 540-720ms (pclust =0.0028; peak at 640ms: M=0.534±0.042). As for what concerns vowels, the two alternatives in our design (/i/ and /o/) differ in both height and backness, precluding the isolation of phonetic sub-classes. Nonetheless, Figure 2C shows that vowel identity was reliably discerned in between 260 and 600ms (pclust =0.0001; peak at 480ms: M=0.596±0.08, chance=0.5) and from 760ms onwards (pclust =0.0001; peak at 860ms: M=0.56±0.067, chance=0.5).

**Figure 2:**
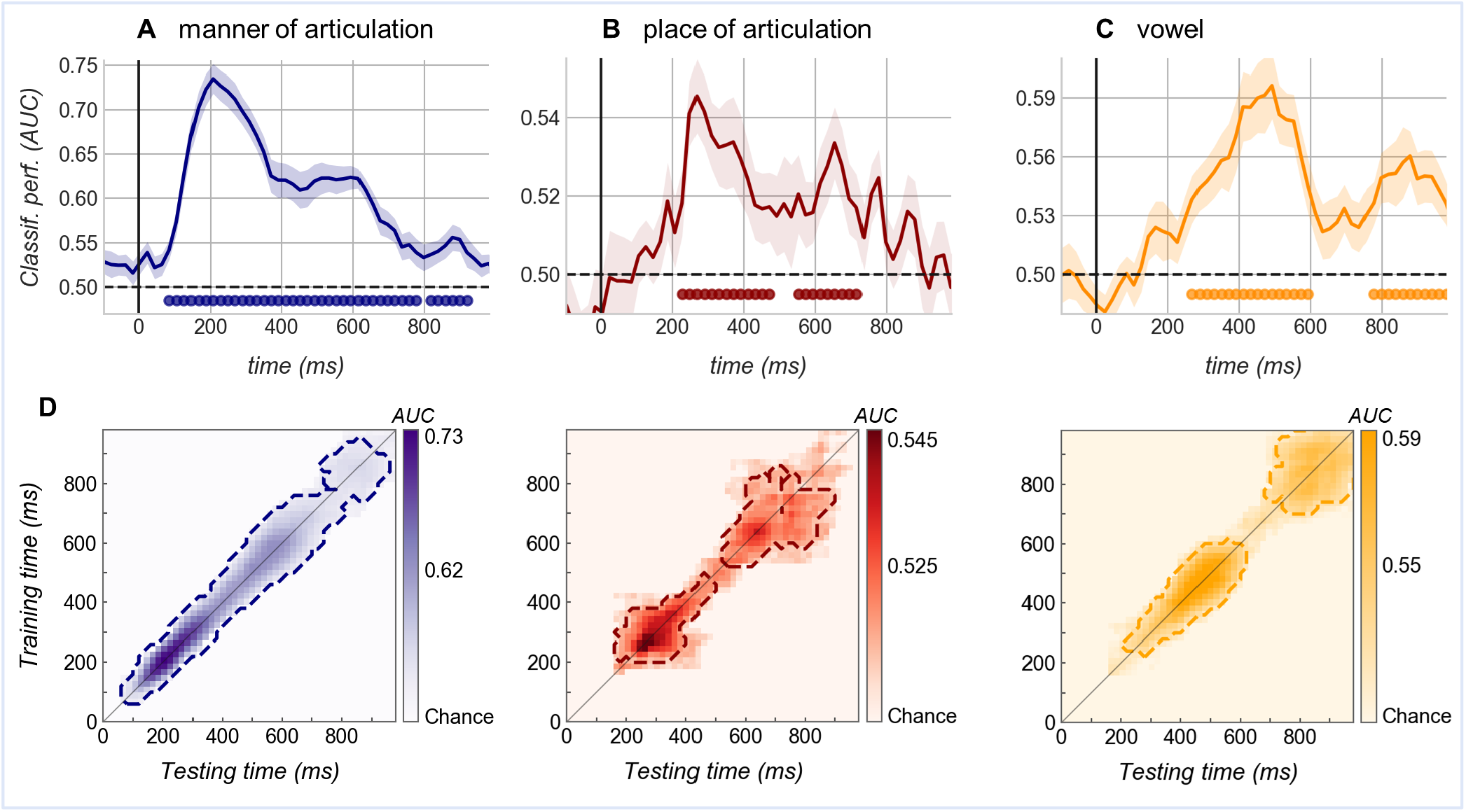
Classification performances of estimators trained on single time windows (20ms) along the ERP. Top: Estimators are tested at the trained time sample. Shaded areas correspond to the standard error (SEM) across subjects, dotted black lines mark theoretical chance level and filled circles indicate significant scores (cluster-corrected t-test). (A) Performance of classifiers trained on manner distinctions: obstruents (/b/, /d/, /g/) vs. sonorants (/m/, /n/, / ɲ/). (B) Performance of classifiers trained on place distinctions: labials (/b/, /m/) vs. alveolars (/d/, /n/) vs. velars (/g/, /ɲ/). (C) Classification of vowel identities: /i/ vs /o/. (D) Temporal generalization matrices: each panel displays above-chance decoding scores of estimators trained on a single time window (y-axis) and tested at every possible time sample (x-axis) along the ERP. The diagonal thin lines demark classifiers trained and tested on the same time sample. Dashed contours indicate significant clusters (manner: p_clust_ =0.0001; place: p_clust_ =0.0001 and 0.0028, vowel: p_clust_ =0.0097 and p_clust_ =0.0108).

To fully characterize the neural dynamics underlying such performances, the same classifiers were systematically tested on their ability to decode across time. In case a neural activation is maintained or recursive, a successful estimator (which is specific to a certain pattern of brain activity) will achieve above-chance scores at multiple time points (King & Dehaene, 2014). Figure 2D illustrates how classifiers generalized only over a limited amount of time lags, indication that the neural activity was progressing along a functional pathway. Concretely, the “cone” shape arising from the generalization matrices discloses the retrieval of evolving neural codes: the activity supporting classification was either transferring across cortical regions, transformed within the same region over time or both. Presumably, the mild widening of the generalization performance observable in the second portion of the trial might denote a change in the representational format reached relatively late after syllable onset.

We started to objectivize these interpretations by using classifier weights to reconstruct informative activity patterns (see Methods). Discriminative activity was diffuse over the scalp, resembling the auditory ERP topographies arising from multiple perisylvian sources that are typical of this age (Figure S3). Crucially, substantiating the occurrence of distinct encoding stages, informative clusters were qualitatively different during the first and second time-windows that provided reliable classification. Change was particularly appreciable in the individual topographies (Figure S3A-B) which are free of the blurring effect created by averaging across participants. We additionally observed that sensors supporting manner and place classification were somewhat separable (Figure S3); and found significant differences between brain activity patterns precisely distinctive for either labials, alveolars or velars (Figure S4, where a detailed overview of place-informative activations is also reported). These findings uncover that infant syllable perception is supported by discrete and local, although distributed and partially overlapping, neural responses, as described for adults (E. F. Chang et al., 2010; Correia et al., 2015).

### A stable code across speakers and co-articulated components

Second, we examined the invariance of the neural code by training new sets of manner and place estimators on a single context (e.g. stimuli spoken by the female voice) and testing them on the alternative untrained condition (e.g. stimuli pronounced by the male voice). We considered the speaker context in a first analysis and the co-articulation in a second analysis. Since several adult and infant studies have shown that linguistic and non-linguistic information are encoded separately from an early processing stage (Formisano et al., 2008; Bristow et al., 2008), we expected full generalization across voice genders. Concerning the co-articulatory context, either infants computed only holistic syllable representations - in that case no generalization should be observed (e.g. an estimator that discriminates “bo” versus “do” on the basis of a whole-syllable code would perform at chance when tested on “bi” versus “di”) - or consonants are encoded independently of the subsequent vowel, leading to successful cross-condition classification.

For manner, the timing of cross-context decoding was virtually identical to that seen in the overall analysis, and the accuracy only marginally reduced (Figure 3A; Tables S1 and S2). Such generalization proves that the infant brain encodes manner features uniformly and irrespective of harmonic particularities, corroborating and extending previous behavioral evidence from older infants (Hillenbrand, James, 1983). Remarkably, clear generalization across voices and vowels was obtained also for place (Figure 3B). The time-course of classification, with two distinct decodable periods, and its accuracy were comparable to those achieved in the initial analysis (Figure 3B, Tables S1 and S2). Since the acoustic cues for place vary considerably with the context (Liberman et al., 1967; Dorman et al., 1977), these cross-condition performances clearly reveal that the infant brain is able to extract an invariant phonetic code beyond acoustic differences, even in the challenging case of place contrasts.

**Figure 3:**
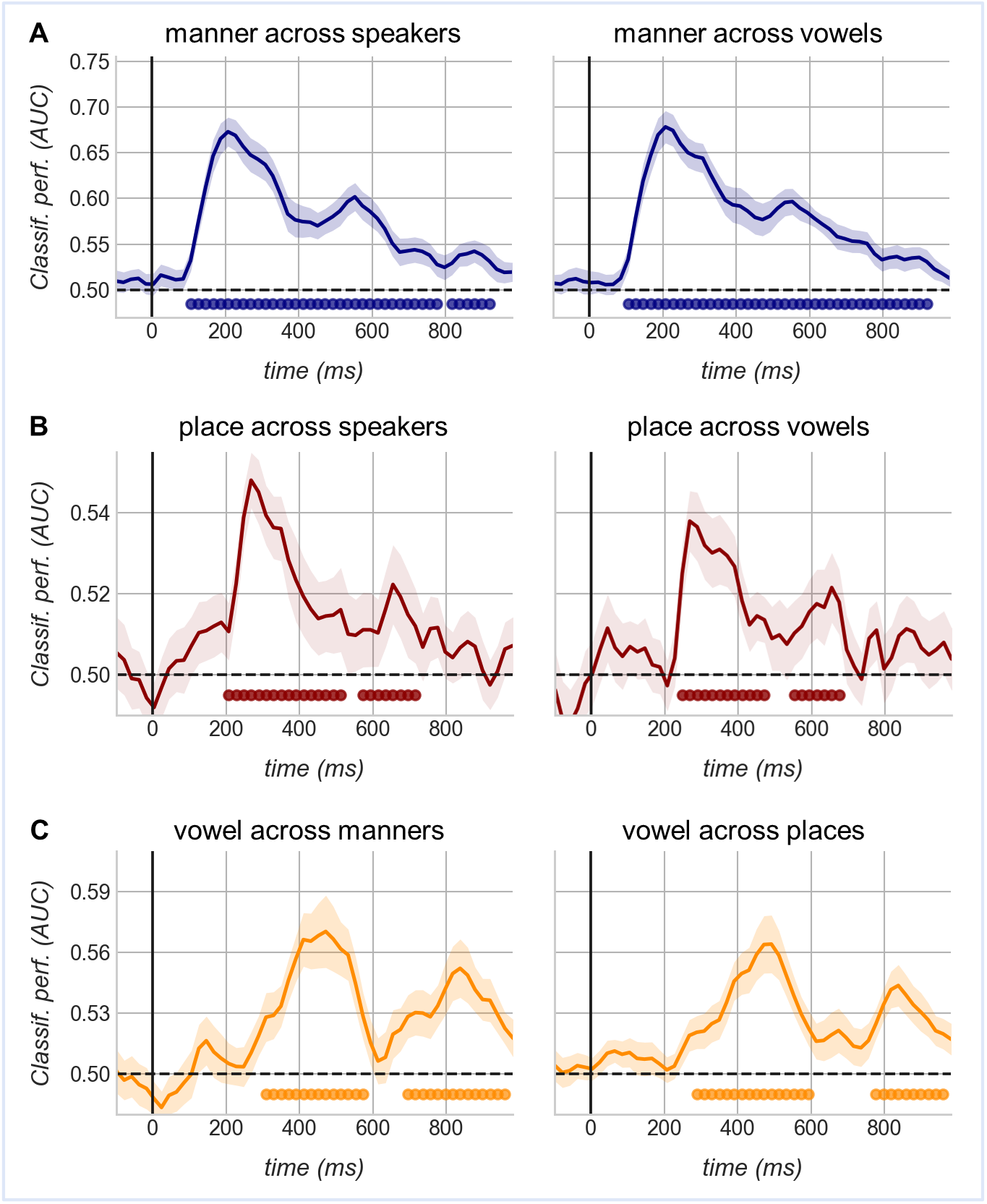
Cross-condition decoding. (A) Left: generalization of manner estimators across voice conditions: classifiers trained on syllables produced by one speaker are tested on stimuli uttered by the other speaker. Right: generalization of manner estimators across vowel conditions: classifiers trained on consonants associated to one vowel are tested on syllables containing the alternative vowel. (B) Same as A, but for place estimators. (C) Left - vowel classification across manners: classifiers are trained on obstruents then tested on sonorants and vice versa. Right - vowel classification across places: vowel estimators are trained on one place condition (e.g. labials) and tested on the other two (e.g. alveolars and velars). Shaded areas correspond to the standard error (SEM) across subjects; dotted black lines mark theoretical chance level. Filled circles indicate scores significantly above-chance (exact p-values are reported in Table S1). Performances from all possible training/test directions are averaged.

Complementarily to these results, vowel estimators trained on single manner or place conditions fully generalized to the alternative contexts (Figure 3C and Table S1). Thus, the cross-decoding patterns observed so far demonstrate that syllables are not perceived holistically but are broken down into sub-components independently of the coarticulated vowel for consonants, and consonantal features for vowels.

### Syllables are factorized into phonetic features, which are secondarily integrated into consonant codes

Note that holistic and unrelated representations of each of the six consonants might suffice for classifiers to sort trials in arbitrary subsets (e.g. /b/,/d/,/g/ vs /m/,/n/,/ɲ/), as done in the previous sections. Crucially, if the infant code for speech is truly based on phonetic features, successful classification should be obtained for one featural dimension regardless of the variation in the other phonetic domains. That is to say, estimators would retrieve the same manner code across labials, velars and alveolars and the same place code in obstruents as in sonorants. To evaluate this possibility, we trained sets of estimators at one featural context (e.g. manner classifiers were trained only on labials) and tested them *within* the same (labials) and *across* untrained phonetic contexts (e.g. alveolars or velars). If the code was based on invariant features, the two tests should yield similar performances.

This criterion revealed two distinct stages (Figure 4A): during an early time-window, both manner and place estimators achieved successful generalization, with a classification accuracy approaching that obtained within the trained condition. Initial phonetic representations are therefore based on an orthogonal code for phonetic features. Beyond ~450ms however, classification performance was significantly lower across featural domains as compared to within, suggesting a change in the representational format. Particularly, cross-condition decoding was impossible for place, while manner information was more resilient but nevertheless altered by the variation in place context (Figure 4A). We hypothesized that secondary processing stages encompass the combination of multiple elementary dimensions, i.e. during this later time window features might be merged into a broader code.

**Figure 4:**
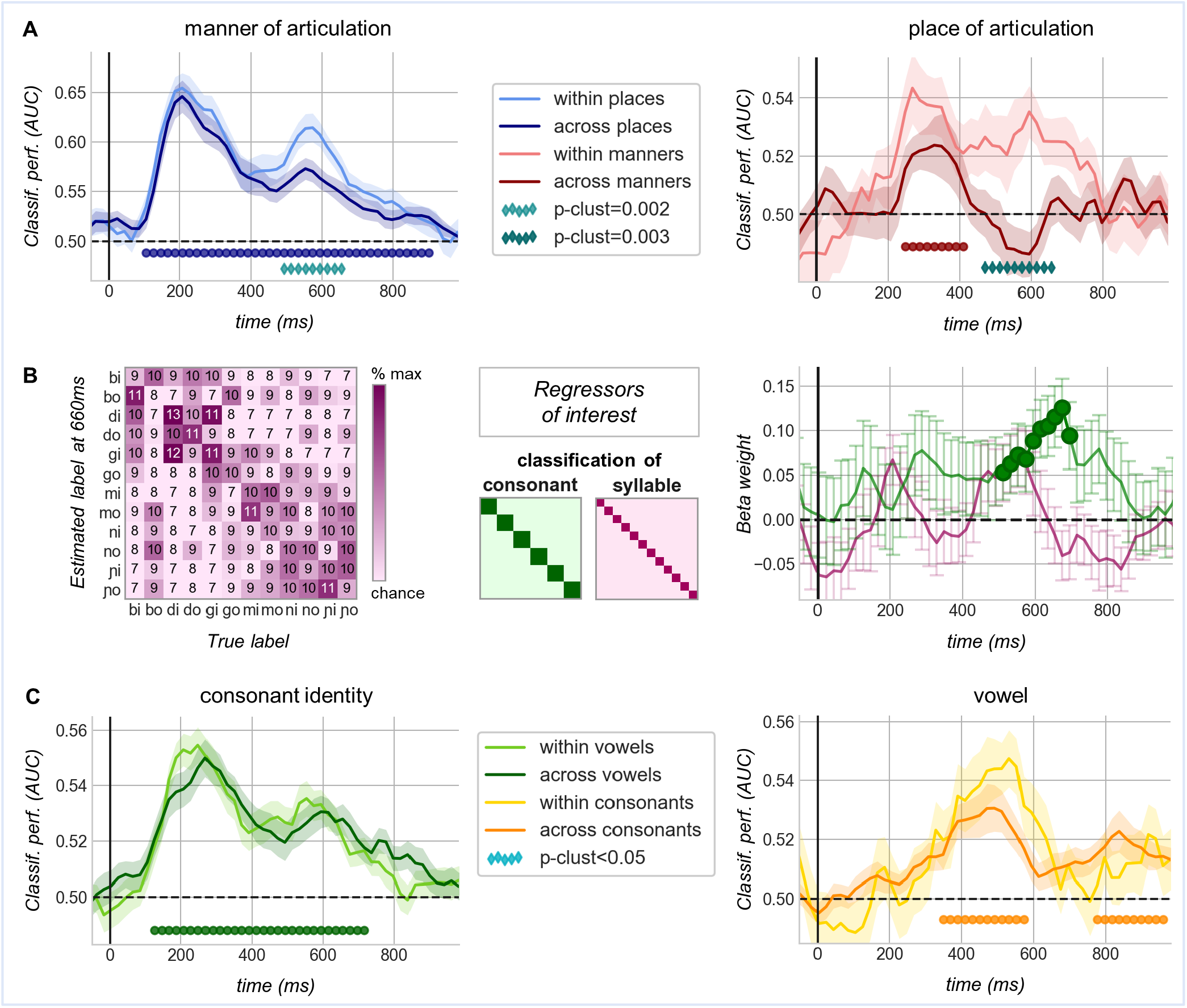
Orthogonal feature codes are merged into phoneme identities at a late stage of processing. (A) Time-resolved performance of estimators trained on a single phonetic feature (e.g. manner estimators trained on labials: /b/ vs. /m/). In light colors: classification within the trained condition (e.g. test on labials); in darker colors: performance at novel phonetic contexts (e.g. test on alveolars: /d/ vs. /n/, and velars: /g/ vs. /ɲ/). Scores from all possible training conditions or train/test directions are averaged. Shaded areas correspond to the SEM across subjects. Filled circles indicate significant generalization across contexts (100-900ms: p_clust_=0.0001 for manner; 240- 420ms: p_clust_=0.001 for place). Diamonds indicate higher performance *within* as compared to *across* conditions (exact time window of significance for manner: 480-640ms; for place: 460-660ms). (B) Left: example of a neural confusion matrix at time t (660ms) obtained with a 12-class (syllables) decoding problem (average across subjects). Numbers within each cell indicate the percentage of times a given syllable from the x-axis was classified with the label reported on the y-axis. Off-diagonal values diverging from 0 signal misidentification (chance=8.3%). Middle: theoretical confusion matrices depicting a perfect separation between (i.e. the ideal classification of) consonants and broad syllable identities (classes are ordered as in the left matrix). Darker colors correspond to the values 50% and 100% respectively, whereas light colors correspond to 0%. These matrices were entered as predictors of interest in a multiple regression analysis to explain neural syllable confusion at each time point. Right: the obtained beta-weights averaged across subjects and marked by filled circles when significantly above zero (cluster-based permutation t-test). Vertical lines correspond to SEM. Three additional predictors (place, manner and vowel discrimination) were entered in the multiple regression as variables of non-interest, their visualization and beta-weights are illustrated in Figure S5. (C) Left: performance of estimators trained on discriminating all consonants (/b/ vs /d/ vs /g/ vs /m/ vs /n/ vs /ɲ/) coupled with one vowel (e.g. “-i”) and tested within the same (light green) and across the other vocalic context (e.g. “-o”; dark green). Right: performance of vowel classifiers trained on a single consonant (e.g. /b/) and tested within the same consonant (yellow) and across the remaining five (orange). Filled circles mark significant generalization across contexts (consonant classifiers: 80-900ms, p_clust_=0.0001; vowel classifiers: 340-560ms, p_clust_=0.0001 and 760ms onwards, p_clust_=0.0002).

To gain additional evidence on a secondary integration stage, we trained algorithms on whole syllable identities (i.e. 12 labels: “bi” vs “bo” vs “di” vs “do” vs “gi” vs “go” etc.) and explored their error patterns at test. In this analysis above-chance accuracy scores (Figure S5A) are difficult to interpret per-se, as class separation might be driven by either one or a mixture of stimuli sub-components. Between-class confusion, on the other end, provides exhaustive information about all the facets of the stimuli encoded by the brain at a given moment. For instance, whereas neural codes based on the whole syllable would produce a purely diagonal confusion matrix, representations based on the identity of the phonemes (i.e. idiosyncratic combinations of manner and place features) would trigger conspicuous mislabeling among pairs of stimuli sharing the same consonant. Using multiple linear regression, we tested whether and when pairwise neural syllable confusion (Figure 4B-left and S5A-bottom) was explained by either consonant and/or whole-syllable codes (Figure 4B-middle) once manner, place and vowel distinctions were entered as variables of non-interest (Figure S5B-top). Complementarily with the decoding outcomes in Figure 4A, the consonant regressor did significantly predict the patterns of neural separability, but *only* in between 500 and 700ms (Figure 4B-right; p_clust_ = 0.006). Conversely, the syllable regressor never reached significance (Figure 4B). Thus, following the encoding of orthogonal features, place and manner codes were integrated into comprehensive consonant representations.

### Consonant and vowels remain separate

Lastly, we queried a possible interconnection between consonant and vowel processing. The results obtained so far contain a few interesting hints in this regard. As shown in Figures 2 and 3, vowel decodability follows a double-peak pattern very similar to that observed for consonantal dimensions, but peak scores are achieved markedly later and at times when consonantal place is hardly discriminable. Together with the invariance of vowel codes across consonantal features (Figure 3C), these observations reveal that infants encoded the two phonemes composing the stimulus orderly and individually.

As a final step, we tested whether the two phonemes were merged into a syllabic unit. Using a similar logic as above, we compared the performance of consonant and vowel estimators *within* and *across* vowel and consonant conditions. The presence of an integrated syllabic code would generate a drop in performance across context. As displayed in Figure 4C, no interaction was found, suggesting that consonant and vowel representations were kept separated, at least until 1 second after syllable onset.

## DISCUSSION

Altogether, the classification patterns observed in this study reveal two speech encoding formats in the infant brain. During a first stage each articulatory-phonetic domain is encoded independently, thus each speech instance is characterized by its coordinates along the manner and place dimensions described by linguists. In a second stage multiple features are combined into a unified and idiosyncratic representation, still allowing phoneme classification but hindering full generalization of featural decoding across phonemes. This functional progression is consistent with the dynamic nature of the neural codes as revealed by the matrices in Figure 2D and the corresponding informative activity patterns in Figures S3-4. Lastly, although our experiment was mainly focused on consonant encoding, similar processing stages for vowels are likely.

The present study draws a striking parallelism between the neural underpinnings of preverbal and adult speech perception (e.g. Mesgarani et al., 2014; E. F. Chang et al., 2010; Correia et al., 2015). In addition, it demonstrates that human neural codes for speech are stable even for those phonetic features reflected by inconsistent acoustic correlates (i.e. the place of articulation). While the recovery of articulatory motor patterns has been proposed as a solution to overcome signal variability, we show that a phonetic code is already in place ~12 weeks after birth, when production skills are still severely limited (Kuhl & Meltzoff, 1996). Our results are thus in disagreement with mainstream views postulating that motor patterns acquisition, with the consequent mapping between articulatory movements and acoustic outcomes, mediates a switch from domain-general to language-specific processing (Kuhl et al., 2008; Westermann & Reck Miranda, 2004; Laurent et al., 2017; Vilain et al., 2019). Instead, our findings provide a new representational solution able to account for the speech abilities observed in early infancy: a vectorized encoding system which projects the signal onto a reduced number of relevant orthogonal axes.

The innate, or acquired, origin of this code needs further study to be understood. Broadly speaking, previous neuroimaging research describing sonority-related phonological biases in newborns (Gomez et al., 2014) proves the plausibility of innate linguistic constrains for humans. Strikingly, preterms born at 30 weeks of gestation have been shown to detect place of articulation changes through a network of temporal and frontal brain areas similar to that recruited at later ages in analogous settings (Mahmoudzadeh et al., 2013, 2016). Considering that the vocal production of these neonates is nonexistent and their sucking behavior very weak and poorly organized, this evidence corroborates our claim for a decorrelation between phonetic perception and articulatory-motor skills. Moreover, the fact that the same discrimination abilities are carried by similar cortical regions across different ages points to a continuity in the codes these regions use, and therefore to a genetically determined mechanism. Nevertheless, it has been recently proposed that early orofacial stereotypies such as tongue protrusion/retraction may provide fetuses and newborns with a primordial knowledge of the shape and configurability of the upper vocal tract (Choi et al., 2017). Such information, combined with sound exposure, might foster an integrative/multi-modal representational space for speech before the onset of canonical babbling.

Whichever its origin, the early availability of a code based on phonetic features could play a crucial role in word learning. To discover words, infants must cope not only with acoustical but also with phonological variation due to the segmental context: for example, in order to apprehend that “wet shoes” and “we[p] pants” share the same word “wet”, English infants should apply a rule stating that an alveolar stop consonant borrows the place of articulation from the subsequent stop (Darcy et al., 2009). Phonotactic rules of this sort pertains phonetic features rather than holistic phonemes. Several behavioral studies reported that infants are sensitive to phonotactic cues already by the age of 9 months: they prefer to listen to sequences that are phonotactically legal in their native language (Friederici & Wessels, 1993; P. W. Jusczyk et al., 1993) and use their phonotactic knowledge to find word boundaries in continuous speech (Mattys & Jusczyk, 2001). At this age, coherently with our argument, phonotactic rules are easily learned when expressed at the level of phonetic features while they are not detected when they concern the identity of the phonemes (Saffran & Thiessen, 2003). A featural encoding of speech is further consistent with the documented ability of young infants to use phonetic details in word-referent mapping (Swingley & Aslin, 2002; Ballem & Plunkett, 2005; Fennell & Waxman, 2010). Phonetic features might then correspond to an essential and quickly available building block for human language acquisition.

The present study further demonstrates that manner and place encoding is followed by a combinatorial process, which is still exquisitely phonetic in nature. In this regard, it is worth to highlight that, once taken phoneme identity into account, we found no evidence for a broader, comprehensive syllabic representation (Figure 4B-C). Such (null) result is inconsistent with studies depicting the syllable as the natural unit of speech perception/processing (Räsänen et al., 2018). For example, 4 days after birth neonates can categorize utterances using the number of their syllabic constituents but not the number of phonemes (Bijeljac-Babic et al., 1993). Around one month of age they discriminate changes within well-formed syllables (CVC and VCCCV; C=consonant, V=vowel) but fail to discern the same kind of alteration within chains of consonants (Bertoncini & Mehler, 1981). Besides, pre-school children and illiterate adults access and manipulate syllables far more easily than phonemes (Morais et al., 1986). Whereas these findings have led various authors to designate the syllable as the basis for speech representation, our results refute the hypothesis of a syllabic unit perceived as a whole and thereafter decomposed into phonetic sub-parts. The discrepancy might come from studying online processing, as we did in the current paper, vs. assessing the content of a memory slot, as done in the behavioral studies. Indeed, behavioral paradigms imply the memorization of a given element that is later compared to a new one or manipulated upon request. Only a full articulatory event (e.g. a spoken syllable) or an external pointer to syllable subparts (e.g. a grapheme) might be storable in working memory. Ultimately, our evidence for phoneme-identity neural codes complements adult data (Zhang et al., 2016) in corroborating the reality of the phoneme as psycholinguistic object (Kazanina et al., 2018). Moreover, a neural separation between consonants and vowels is particularly meaningful in light of the proposal suggesting diverging functional roles for these components in language: while consonants are more informative for lexical distinctions, vowels are particularly apt to mark structural organization (Nespor et al., 2003). Coherently with our findings, and just as adults (Toro et al., 2008), infants have been shown to exploit such “division of labor” in order to extract lexical and syntactic information already by the age of 12 months (Hochmann et al., 2011).

In summary, our results indicate that infants project the high-dimensional speech signal onto several axes of neural responsivity corresponding to phonetic features. This process creates a structured and highly generalizable space that is robust to surface variability across speakers or co-articulatory contexts. As outlined for faces (L. Chang & Tsao, 2017), a factorized representational system is more efficient and more flexible than exemplar coding (e.g. Port, 2007; 2010) and therefore ideally suited for the bootstrapping of language acquisition. In support of this claim it has been shown that when non-pertinent acoustic variability is high within the experimental setting (as it is in real-life scenarios) infants are particularly prone to use minimal phonetic contrasts to learn words (Rost & McMurray, 2009). Efficiency and flexibility characterize the second-stage integrative code as well: elementary components, i.e. the phonetic features, are recombined into intermediate representations, i.e. consonants and vowels, optimizing in this way the accessibility of lexicon and syntax (Hochmann et al., 2011).

To conclude, pending more definitive experimental evidence, we point out the possibility that an abstract phonetic code might be available from birth and endow infants with the ability to discriminate phonemes from most languages (Peter W. Jusczyk, 2000). As an additional conjecture, we envision that the second-stage integrative process could be subject to learning: areas downline of the first processing phase might become selective for the most frequently encountered feature combinations. Further experiments, spanning a range of languages and ages, will be needed to investigate how the observed codes adapt to the inventory of native phonemes.

## MATERIAL AND METHODS

### Participants

25 full-term, normal-hearing infants (12 females, 13 males) coming from a French-speaking environment were tested between 12 and 14 weeks after birth (mean age= 12 weeks and 6 days). An additional 16 participants were excluded from analysis because of: excessive agitation during the experimental session (n=6), insufficient number of trials after artifact rejection (n=3, the artifact rejection procedure is described below), technical problems during data collection (n=3), aberrant global field power (GFP) in the average of all syllable-related potentials (i.e. peak GFP<4μV, n=4). The protocol was approved by the regional ethical committee for biomedical research (CPP Region Centre Ouest 1). Parents gave their written informed consent before starting the experiment.

### Stimuli

Stimuli consisted of 120 speech sounds constructed upon 6 consonants: /b/, /d/, /g/, /m/, /n/, /ɲ/. These consonants were selected to cover two manner features, i.e. obstruent (/b/, /d/, /g/) and sonorant (/m/, /n/, /ɲ/), and three places of articulation, i.e. labial (/b/, /m/), alveolar (/d/, /n/), and velar-palatal (/g/, /ɲ/). In case each consonant was spoken always in the same way throughout the experiment, there would have been one-to-one correspondence between the articulatory profiles (e.g. obstruent + labial) of the stimuli and their spectrograms; while our goal was precisely to disentangle phonetic from merely acoustic stimuli representations. Each consonant was therefore associated with two vowels, /i/ and /o/, and produced by a male and female speaker to obtain 2 manner *x* 3 place *x* 2 vowel *x* 2 voice factor design (i.e. 24 sub-conditions). To increase acoustic variability (and extend the external validity of our measurements), speakers were asked to repeat the same tokens several times changing their intonation. For every sub-condition we selected 5 utterances, distinct in low-level acoustic characteristics such as pitch and duration. In the resulting set of syllables each manner of articulation condition contained 60 spectrotemporal profiles (3 consonants *x* 2 vowels *x* 2 voices *x* 5 utterances); similarly, each place of articulation was presented in 40 (2 consonants *x* 2 vowels *x* 2 voices *x* 5 utterances) spectrotemporal versions.

Speech signals were recorded in a silent chamber using a dynamic microphone (Beyerdynamic DT 290 broadcast headset) on a linear PCM recorder (DR-05, TASCAM) at a sampling rate of 44.1 kHz. Recordings were first cleared from background noise in Audacity 2.1.3 (https://www.audacityteam.org) and further edited with PRAAT software (Boersma & Weenink, 2017). Acoustic transients (clicks) were manually removed and stimuli length was adjusted to fall within the range of 350-425ms. Tokens were normalized for peak amplitude and average (i.e. root-mean square) intensity, obtaining maximal audibility and loudness equalization. All stimuli were placed on the left channel and a click was positioned on the right channel at the exact time-point of syllable onset. The left channel was connected to the audio amplifier (mono input to the loudspeakers) while the right channel was connected to the EEG amplifiers through the DIN port to create a TTL signal. Brain voltage and clicks were recorded simultaneously with the same temporal resolution providing a precise mapping between EEG recording and stimulation.

Articulation, and in particular the manner, is known to affect consonant duration, introducing the risk of possible confounds between this low-level cue and the phonetic feature. To validate our set of syllabic stimuli, we therefore assessed consonant lengths through a gating procedure (Grosjean, 1996). Over multiple trials, each stimulus was listened in portions of progressively increasing duration (10ms steps), starting from the end of the syllable and proceeding backwards, toward its beginning. The duration of the longest portion for which no consonantal sound was perceived was subtracted from the total length of the stimulus. Consonant duration assessed in this way ranged between 80 and 210 ms (M±SD=154±25) and varied homogeneously across categories (i.e. /b/, /d/, /g/, /m/, /n/, /ɲ/; F(5,114)=1.42, p=0.222). Most importantly, consonant duration did not change as a function of manner nor place of articulation. In an ANOVA with these two factors, the effect of manner (F(1,114)<1), the effect of place (F(2,114)=1.28, p=0.280) and their interaction (F(2,114)=2.25, p=0.109) were not significant.

### Procedure

Subjects were tested in a soundproof Faraday cage equipped with a computer screen and loudspeakers on the top. Infants were hold by a caregiver, their position was chosen to guarantee personal comfort and at the same time enable good-quality data acquisition. Syllables were broadcast through the loudspeakers at 70 decibels, in a latin-square randomized order and with an inter-stimulus interval (ISI) randomly picked between 600 and 1000ms. To minimize body movements we presented engaging visual animations that were unsynchronized with the auditory stream. Sleep was highly encouraged at any time; on average our subjects slept for 65% of the experimental session. Pauses were made whenever needed. The experiment finished with the presentations of 3136 tokens (corresponding to approximately 63 minutes of listening time) or as soon as infants became restless.

### EEG recording and data preprocessing

The electroencephalogram (EEG) was continuously digitized at 500 Hz (Net Amps 300 EGI amplifier combined with NetStation 5.3 software) from 256 channels. We used a prototype HydroCel net (EGI; Eugene, OR, USA) referenced to the vertex. The sensor layout of this prototype diverges from the classical geodesic 128-locations partitioning (Tucker, 1993) in that 20 of the standard temporal positions are covered by 2 tight grids of sensors (70 electrodes on each side, organized in hexagonal pods) with no sponge inserts (Figure S2). Electrodes are made of carbon fibers embedded within a plastic (ABS) substrate and coated with silver-chloride.

### Artifact detection and correction

Data preprocessing was conducted through custom-made MATLAB scripts based on the EEGLAB toolbox 14.0 (Delorme & Makeig, 2004). While following the main preprocessing steps normally used in developmental studies, we introduced some modifications inspired by efforts carried to improve adult data quality (Jas et al., 2017; Mognon et al., 2011). Namely, we identified artifacts on the continuous recordings with the employment of adaptive rather than absolute/predefined thresholds. In this way, we could account for inter-individual variability and the heterogeneous influence that reference distance and vigilance state exert on the voltage. Moreover, we did not discard but corrected local and transient artifacts, exploiting the redundancy of information provided by our dense sensor-layout (Figure S2) and high sampling rate.

As a first step, EEG recordings were band-pass filtered ([0.5 - 40Hz]) and the mean voltage of each electrode was set to zero. Artifacts were detected before segmentation by a series of algorithms with adaptive thresholds. These algorithms rejected samples on the basis of: the voltage amplitude and its first derivative; the variance across a 500ms-long moving time window; the fast running average and the deviation between the fast and the slow running averages within a 500ms-long sliding time window. Thresholds were set independently for each subject and for each electrode upon the distribution of these measures along the whole recording (threshold = median +/- *n*\*IQ, where IQ is the interquartile range of the distribution). Two additional algorithms identified whether the power within the 0-10Hz band was excessively low or within 20-40Hz excessively high relative to the total power; and whether the voltage amplitude displayed by each sensor at a given time point was disproportionate relative to that recorded by the other sensors at the same instant. For these last two algorithms, thresholds were computed upon the distribution across channels.

The output of the artifact detection procedure was a rejection matrix with the same size of the EEG recording. We used this matrix to mark time points with prominent artifacts *(bad times)* and channels that did not function properly *(bad channels).* We identified as *bad times* periods longer than 50ms with a percentage of rejected channels superior to 30% or beyond 2IQ from the 3^rd^ quartile of the distribution of the percentage of rejected channels across time. Similarly, *bad channels* were the ones not working properly for more than 30% of time or with a percentage of bad samples that went beyond 2IQ from the 3^rd^ quartile of the distribution of the percentage of rejected samples across channels.

Periods defined as *bad times* were not corrected because there was not enough information available to reconstruct the signal. For the rest, two kinds of correction were applied. When the rejected segments had a very short duration (50ms max, e.g. heart beats or jumps) we relied on the assumption that, during these periods, most of the variance came from noise. For each of them, principal components were estimated (PCA) and the first *n* components determining 90% of the variance were removed. Otherwise, we corrected *bad channels* and long rejected segments that did not contain *bad times* using spherical splines interpolation (Perrin et al., 1989). Spatial interpolation was carried out only if at least 50% of the neighboring channels were intact. Corrected segments were realigned with the rest of the data which were then high-pass filtered (0.5Hz) to eliminate possible drifts resulting from this operation. The artifact detection-correction procedure was applied iteratively, keeping previously identified bad samples aside for the subsequent artifact detection steps.

### Epoching

EEG recordings (and the corresponding rejection matrix) were segmented into epochs starting 200ms before and ending 1400ms after syllable onset. Trials were rejected if more than 15% of their samples contained artifacts. Epochs were also discarded based on their Euclidean distance from the average, i.e. when their mean or maximum distance from the average response was an outlier in the distribution (> 3^rd^ quartile + 1.5*IQ). Following automated rejection, the remaining epochs were visually inspected and a few trials still presenting obvious aberrancies were manually eliminated.

Since multivariate pattern analysis requires a conspicuous amount of trials, we included subjects with a minimum of 40 trials/sub-condition. In our final group of infants (N=25), the mean trial rejection rate was 28.7% (12.4 to 53.5%). On average, the number of artifact-free epochs available per subject in each sub-condition (e.g. “bi-female”) was 70, providing 840 trials for each manner of articulation condition and 560 trials for every place of articulation condition.

Before submitting them to the main analyses, epochs were low-pass filtered at 20Hz, mathematically re-referenced to the mean of all channels and down-sampled (with a moving average of 2 time points) to 250Hz. All the main analyses (decoding) were carried at the single trial level. Nonetheless, epochs were also averaged per either subcondition or manner-/place-condition in order to examine evoked responses (ERPs, e.g. Figure S4C).

### Decoding

Multivariate pattern analyses were conducted within subject, relying on the Scikit-Learn (Pedregosa et al., 2011) and MNE (Gramfort et al., 2013, 2014) Python packages. To decode *in time* epochs were divided into 300 consecutive windows of 20ms (from −200ms to 1000ms relative to stimulus onset), each corresponding to a matrix with the shape *n* channels *x* 5 samples (sampling rate = 250Hz, 5 samples=20ms). Each analysis was carried on a single window with the general aim of predicting a vector of categorical data (*y*) from a matrix of single-trial neural data (*X*) which included all EEG channels. To decode the manner of articulation trials were labelled as belonging to either the category of “obstruent” or to the category of “sonorant” depending on whether /b/, /d/, /g/ or /m/, /n/, /ɲ/ exemplars were presented. To decode the place of articulation *y* comprised three classes: “labial” (/b/, /m/), “alveolar” (/d/, /n/), and “velar” (/g/ and /ɲ/). For vowel decoding, trials were separated in two classes, “i” and “o”, based on the vocalic portion of the stimulus.

All decoding analyses were performed within a stratified cross-validation procedure consisting of 100 iterations. At each run, trials were shuffled and then split into a training and a test set containing 90% and 10% of trials respectively. As compared to the most common folding approach, this cross-validation outline enabled to maximize the number of iterations (and thus the reliability of the final performance) while maintaining a fixed and reasonable amount of test trials. Importantly, stratification ensured (a) that the same proportion of each class was preserved within each set (b) all sources of variability (e.g. voice gender) were evenly represented across sets (e.g. training and test sets contained syllables produced by the female vs male speaker in the same proportion).

Given the high amplitude fluctuations typically seen in infant EEG background activity, we first aimed at improving our signal-to-noise ratio. Once defined the training and the test set for a given run, we applied a “micro-averaging” procedure, a strategy previously used on adults with the same purpose (Grootswagers et al., 2016). This consisted in averaging together randomly picked groups of 16 epochs within each class. The number of trials to average being arbitrary, we tried with 4, 8, and 12 and observed that by averaging 16 trials we could reach the best performance without compromising its reliability. Note that such assessment was conducted on the first decoding analysis we had planned (i.e. manner of articulation within a standard cross-validation schema) and the choice of 16 was then adopted a priori for all the other decoding analyses. At the end of this operation, to ensure perfect balance among classes, we equalized the number of (micro-averaged) epochs across categories. In practice, this consisted in dropping 1 to 3 randomly picked trials from the most numerous class(es).

Next, following the z-scoring each feature (i.e. channel and time point across trials), a L1-norm regularized Logistic Regression (Fan et al., 2008) was fitted to the training set in order to find the hyperplane that could maximally predict *y* from *X* while minimizing a log loss function. L1 penalty was chosen to exclude less informative features from the solution (their weights being set to zero). Such regularization can be conceived in terms of dimensionality reduction, an optimization that enabled us to prevent overfitting (by reducing model complexity (Ng, 2004)) but still exploit the high density of our EEG data. The other model parameters were kept to their default values as provided by the Scikit-learn package. When decoding concerned more than two classes (e.g. place classification) we adopted a “one-vs-rest” approach: for each class (i.e. each place of articulation) one model was fitted against all the other classes.

Once trained, the models were used to predict *y* from the test set and their performance was evaluated by comparing estimates to the ground truth. All algorithms produced as an outcome vectors of probabilistic estimates. These probabilities were scored by computing the area under the Receiver Operating Characteristic curve (AUC), which summarizes the ratio between true positives (e.g. trials correctly classified as “obstruent”) and false positives (e.g. trials classified as “obstruent” while a sonorant consonant was presented). The value of AUC ranges between 0 and 1, with 0.5 corresponding to chance level. Once again, in multiclass decoding a “one-vs-rest” scheme was used: the AUC scores were computed for each class against all the others and then averaged. Lastly, for both binary and multiclass problems, evaluations were averaged over all cross-validation runs.

As a proof of concept, the main decoding analyses were performed with two additional algorithms: L1-norm regularized linear Support Vector Machine (SVM; (Fan et al., 2008)) and Linear Discriminant Analysis (LDA). For the latter, a shrinkage estimator of the covariance matrix was used, taking into account the fact that the dimensionality of our data vectors exceeded the number of samples in each class (Ledoit & Wolf, 2003). Importantly, we restricted our alternatives to linear classifiers to make sure that the algorithms focused on explicit neural codes (Kriegeskorte, 2011). Beside slight variations in accuracy, alternative classifiers yielded very similar outcomes.

### Generalization across time (Figure 2D)

Estimators trained at each time window *t* were systematically tested on (both the same and) every other possible time window *t’,* i.e. every 20ms from 200ms prior to 1000ms after syllable onset. Such procedure was performed within the cross-validation so that training set at *t* and test set at *t*’ came from different groups of trials. In the resulting “temporal generalization matrices” each row corresponds to the time lag at which the estimator was trained and columns correspond to the time windows at which it was tested (King & Dehaene, 2014). The shape of the performance within these matrices provides peculiar insights upon the dynamics of the underlying brain activity. If the same neural code was found at *t* and *t*’, the classifier trained at *t* would generalize at *t’.* If, on the contrary, information was passed to another stage of processing characterized by its own coding scheme, performance at *t*’ would be at chance (King & Dehaene, 2014).

### Generalization across conditions

We examined the consistency of information used by classifiers in different harmonic and co-articulatory contexts by performing cross-condition decoding. To ask whether the same neural codes supported the classification of phonetic features and vowel identities across different harmonic contexts, we trained estimators on manner contrasts (/b/, /d/, /g/ vs /m/, /n/, /ɲ/); place contrasts (/b/, /m/ vs /d/, /n/ vs /g/, /ɲ/) and vowel contrasts (/i/ vs /o/) within one speaker condition (e.g. syllables pronounced by the female voice) and tested these same estimators on the other speaker condition (e.g. syllables spoken by the male voice). The procedure regarding co-articulations was analogous: we trained place and manner estimators on one vowel context and tested them on the other; we trained vowel estimators on single manners or places and assessed their performance on the alternative ones.

To test the orthogonality of manner and place encoding we trained estimators on each featural condition separately. More specifically, to reveal place-independent phonetic processing classifiers were trained on the manner comparison (“obstruent” vs “sonorant”) at single place contexts (e.g. only labial sounds). These estimators were then tested both at the trained place (e.g. labials) and at the two unseen places (e.g. alveolar and velar consonants). In case manner neural codes were independent from the place of articulation, we expected classifier to perform comparably *within* the trained place and *across* unseen place contexts. Following the same rationale, we asked whether place codes are specific to manners of articulation by training classifiers to discriminate labials vs. alveolars vs. velars on one manner (e.g. only with obstruent sounds) and testing them within the same (e.g. obstruents) and at the alternative manner condition (e.g. sonorants).

Moreover, we investigated the orthogonality of consonant and vowel representations with two complementary procedures. First, we trained algorithms to distinguish each consonant based on single vocalic contexts (e.g. separation of /b/ vs /d/ vs /g/ vs /m/ vs /n/ vs /ɲ/ when they were co-articulated with /i/) and tested them within the same and across the alternative co-articulatory context (e.g. classify consonant identity among “bo”, “do”, “go”, “mo”, “no”, “ɲo”; note that for this schema, as for place classification, we adopted a “one-vs-rest” approach and the percentage of correct classifications as evaluative metric). Analogously, we trained vowel classifiers on each consonantal option and assessed their performance within the trained consonant and across the five alternative ones. In case consonant and vowel were represented separately, we expected to obtain comparable scores *within* and *across* conditions; oppositely, a degradation in performance across conditions would be indicative of interdependence between the two.

For cross-condition decoding we modified the cross-validation scheme described above so that models fitted on each training set were directly applied at all trials belonging to the untrained condition (i.e. the test set “*across*”). In this way, we capitalized on the independence of train and test sets. Concerning the splitting of single-condition datasets (i.e. the dataset “*within*”), the number of test trials was calibrated to guarantee a minimum of 2 microaveraged trials/class at test and at the same time maximize the amount of trials available for training. Note also that in order to ensure an adequate number of training/test samples, the micro-averaging for the last two cross-decoding schemas was reduced to groups of 8 epochs. Apart from these modifications, the decoding procedures resembled those described above.

### Weight projection (Figure S3)

The weights assigned by classifiers to EEG sensors reflect the degree to which the information captured by a given sensor is used to maximize class separation. However, weights per se are very difficult to interpret. For example, higher weights do not necessarily correspond to high levels of class-specific information as they could be assigned to sensors that are employed to delineate and suppress noise (for a full explanation see (Haufe et al., 2014)). To overcome this issue it is possible to project weights back onto an interpretable activation space by multiplying them with the covariance in the data (cov(X), where X is the N × M matrix of EEG data with N trials and M channels). In the resulting vector (that has length M channels) large amplitudes indicate high degrees of class-specific brain activity (Grootswagers et al., 2016; Haufe et al., 2014). Since our goal was to reconstruct informative activity peculiar to each phonetic feature domain, we retrieved the coefficients of classifiers trained within each place condition to obtain “pure” manner-distinctive patterns and trained within each manner condition to obtain “pure” place-distinctive patterns. By doing so, we ensured that no information about place was available to manner estimators and no information about manner was available to place estimators. After multiplying coefficients and EEG covariance, the resulting activity estimations were averaged across places (to obtain informative activity for manner) or manners (to obtain informative activity for place).

To identify sensors that were crucial specifically for manner or crucial specifically for place classification, we computed the 10th and 90th percentiles of the informative activity values observed throughout the trial. At each time point, channels whose informative activity amplitude fell below the 10th or above the 90th percentiles in one phonetic domain but not the other were interpreted as particularly important to manner but not place classifiers or vice versa (Figure S3).

### Neural syllable confusion and multiple regression analysis

For this section we first built a twelve-class decoding problem by pulling together the female and male conditions and then training algorithms to separate each syllable from all the others (i.e. “bi” vs “bo” vs “di” vs “do” vs “gi” etc.). We adopted a “one-vs-rest” approach and used the same pre-processing steps described for the main analyses. Within each cross-validation loop, we stored the error matrices displayed by these classifiers at test. After averaging across runs, we obtained a series of matrices where the entry at row *i* and column *j* corresponds to the percentage of samples belonging to class *j* and labeled as *i* by the classifier (Figure 4B-left and S5A-bottom). The diagonal of these confusion matrices depicts class-wise accuracy, with theoretical chance being at 8.3% (Figure S5A-top). Given that there is a variety of stimuli characteristics other than syllable identity which could lead to above-chance scores (up to 50%), diagonal entries alone are hardly interpretable. On the other hand, misclassification patterns (i.e. off-diagonal entries in the matrices) have the potential to reveal which dimensions of the stimuli the neural code honors or disregards. To uncover the neural representational geometry (Kriegeskorte & Kievit, 2013) captured by our algorithms and its evolution over time, we employed multiple linear regression. Specifically, we modeled each confusion matrix as a linear combination of five classification performances: those of the ideal manner, place, vowel, consonant and whole-syllable decoders (Figures S5B-top and 4B-middle). Concerning the matrix modelling manner discrimination, for example, the predicted entries for those pairs of syllables sharing the same manner correspond to 16.6%, whereas the predicted value for pairs of syllables not sharing the same manner is 0%. The five predictors were used to explain the (neural) syllable confusion observed at each time point, generating a vector of beta-weights for each of the five regressors. All matrices were z-transformed before estimating the coefficients. With this multiple regression approach we capitalized on the opportunity to separate the potential impact of new variables of interest (i.e. consonant and holistic syllable, Figure 4B) from that of influential dimensions already isolated by the previous analyses (i.e. manner, place and vowel, Figure S5B) on syllable confusion patterns. Significantly above-zero beta-weights assigned to a particular regressor indicate that, at a given time point, the classifier relies on the dimension reflected by that model over and beyond the remaining four variables.

### Statistical analysis

To calculate statistics we performed second-level tests across subjects employing the MNE dedicated functions. Following the example in (Jean-Rémi King et al., 2016), we tested whether (a) time-resolved classification scores were higher than chance; (b) time-resolved classification scores within the trained context were superior to those across context; (c) whether multiple regression beta-weights were higher than zero; using one-sample cluster-based permutation t-tests (Maris & Oostenveld, 2007) which intrinsically account for multiple comparisons. The analyses considered one-dimensional clusters in all cases apart from the generalization across time matrices (with shape training times *x* testing times) for which clusters were bi-dimensional. Univariate t-values were calculated for every score/beta-weight with the exclusion of those corresponding to the baseline period. All samples exceeding the 95^th^ quantile were then grouped into clusters based on cardinal or diagonal adjacency. Cluster-level test statistics corresponded to the sum of t-values within each cluster. Their significance was computed by means of the Monte-Carlo method: they were compared to a null distribution of test statistics created by drawing 10000 random sign flips of the observed outcomes. A cluster was considered as significant when its p-value was below 0.05.

We compared labial-, alveolar- and velar-specific patterns of informative activity with 1-way repeated measures ANOVA. Similarly to above, we addressed the multiple comparisons problem with a permutation procedure based on spatio-temporal clusters. Neighboring elements that passed a threshold corresponding to a p-value of 0.01 were grouped together and their significance was computed by comparing cluster-level statistics to a null distribution of f-value sums created by drawing 10000 random permutations of the observed data. Again, a cluster was considered as significant when its p-value was below 0.05. Since informative activity patterns are meaningful only in case of successful decoding (Haufe et al., 2014), differences were evaluated only during the two time windows when place classification was reliably above chance.

## ACKNOWLEDGMENTS

This research was supported by grants from the Fondation NrJ, Fondation Bettencourt and from the European Research Council (ERC) under the European Union’s Horizon 2020 research and innovation program (grant agreement No. 695710). We are grateful to Don Tucker and Amy Rowland (EGI and University of Oregon) for designing the 256-electrodes net; and to Bahar Khalighinejad for providing help with auditory spectrogram estimation. We are also thankful to Stanislas Dehaene, Yair Lakretz and Christophe Pallier for constructive feedback and suggestions. This paper is dedicated to Sébastien Marti, whose mentorship was essential. His smile, kindness and friendship remain in our hearts.

## AUTHOR CONTRIBUTIONS

G.D.L. conceived and supervised the project. G.G. implemented the experimental design. M.P. and G.G. collected the data. G.G. performed the analysis. G.G. and G.D.L. wrote the manuscript. A.F. provided the pre-processing tools and manuscript revision.

## Supplemental Information for

### Supplementary text

#### Auditory spectrogram estimation and Representation Similarity Analysis

This preliminary investigation was aimed at delineating the auditory representational geometry elicited by our stimuli set (Kriegeskorte, 2008; Kriegeskorte & Kievit, 2013).

The time-frequency auditory representation of the speech sounds was estimated according to a model of the peripheral auditory system (Chi et al., 2005) as implemented in the NSL Matlab Toolbox (http://nsl.isr.umd.edu/downloads.html). This model comprises: a first step in which sound frequencies are spatially separated along the basilar membrane; a second stage that simulates the transduction of basilar membrane displacements into auditory nerve spikes; and a third phase of processing within the cochlear nucleus. The output of the model is an auditory spectrum of the signal as it enters the inferior colliculi. The three stages and their mathematical implementations are described in (Yang et al., 1992) and (Wang & Shamma, 1994). Auditory spectra were computed based on consecutive windows of 10ms for each stimulus, obtaining a total of 120 bidimensional (time *x* frequency) auditory representations. We then estimated pair-wise auditory dissimilarity following two different approaches.

First, we calculated time-resolved auditory (dis)similarity. For this purpose, spectrograms were aligned upon the consonant offset times determined with the gating procedure described in the Materials and Methods *(Stimuli* section). Consonant offset was preferred over syllable onset because acoustic cues for the place of articulation are generally proposed to reside within the formant transitions (i.e. at the time of the switch between consonant and vowel portions) (Liberman et al., 1954). Since consonant duration varied across speech tokens, alignment based on syllable onset would have led to a jittering of such transition times across spectrograms and this jittering could have misleadingly attenuated relevant cues. The 5 auditory spectrograms corresponding to each sub-condition (e.g. the 5 utterances of “go-female”) were then averaged together (Figure S1B). For each (10ms long) spectral frame, we z-scored amplitude values across frequencies and calculated the Euclidean distance between each pair of subconditions. Standardization was applied in order to maximize our power of detecting phonetic distinctions despite variation in fundamental frequencies (i.e. despite male and female voices being characterized by very distinct pitches). The choice of the Euclidean metric is justified by its potentiality to mimic infant discriminative behavior with higher fidelity relative to other distance measures (Sundara et al., 2018). The outcome of this first approach is a series of 35 auditory distance matrices (Figure S1B), describing all together how pairwise auditory (dis)similarity unfolds over time.

It has been proposed that the acoustic correlates of the place of articulation, a feature of major interest in the current study, have an integrative and dynamic nature (Nossair & Zahorian, 1991). The employment of brief time slices could have then potentially precluded us from capturing meaningful cues derivable from the spectral shape as a whole. To account for this eventuality, our second approach relied on the Dynamic Time Warping (DTW) algorithm (Sakoe & Chiba, 1978; Park & Glass, 2008) as implemented in the Python module *dtaidistance* (Meert & Van Craenendonck, 2018). This technique enabled us to find the best alignment between each pair of spectrograms by stretching and compressing them locally, along the time axis. Following z-scoring, we estimated the DTW distance between each pair of utterances and obtained a comprehensive auditory dissimilarity matrix by averaging the distance values corresponding to each pair of sub-conditions.

To investigate the relationship between the auditory space and the phonetic/harmonic dimensions of our speech stimuli we tested the correlation of the auditory distance matrices with four theoretical matrices (Figure S1C). The latter consisted of categorical models in which two syllables are identical (dissimilarity = 0) if they share the same manner/place/vowel/voice, and different (dissimilarity=1) in case they do not. Concerning place of articulation distinctions, some investigations in phonetics seem to suggest that labials/velars and alveolars could be acoustically closer to each other relative to labial and velars (Cho & Ladefoged, 1999; Lisker & Abramson, 1964). Furthermore it has been proposed that the alveolar feature may be “underspecified” (i.e. coronal may correspond to the default place and therefore be somehow inactive/less contrastive) as compared to the labial or velar features (Cummings et al., 2017; Stemberger & Stoel-gammon, 1991; Tsuji et al., 2015). To account for these possibilities, we built an additional model where the distance between labials and alveolars and that between alveolars and velars was quantified as “0.5”. Results obtained with the two place models were completely overlapping.

The match between auditory and theoretical dissimilarity matrices was quantified with a Mantel test for two-dimensional correlations (Mantel, 1967) employing Spearman’s rho as test statistic and performing 10000 permutations for each test. The Mantel procedure, unlike the classical correlation methods, enabled to account for the fact that distances here were not independent, i.e. every dissimilarity depended on two spectral patterns/qualitative values, each of which also codetermined the similarities of all its other pairings in the matrix. For what concerns the time-resolved outcomes, false discovery rate (FDR) correction was used to control for multiple comparisons across spectral frames and results are show in Figure S1D. The comprehensive auditory dissimilarity matrix was significantly correlated with manner (Mantel *rs* =0.228, p=0.0002); vowel (Mantel *rs* =0.297, p=0.0001) and speaker distinctions (Mantel *rs* =0.24, p=0.0001) but not place of articulation (Mantel *rs* =-0.029, p=0.75).

As a note, the reader may wonder the reason why we could not apply the same decoding strategies used on neural data in order to characterize the auditory space. Generally speaking, the lower the number of samples and the higher the ratio of features to sample size, the more a machine learning model will fit the noise in the data instead of a meaningful underlying pattern (Jain & Chandrasekaran, 1982; Kanal & Chandrasekaran, 1971). In the case of our auditory spectrograms, algorithms would need to be trained/tested on a maximum of 120 samples with 4480 features each (as a benchmark: samples for each neural estimator in the main analyses were approximately 1600 and contained 1260 features each). Evidently, such disproportionate dataset is ill-suited for the same kind of estimators used on the ERPs: instability and overfitting would completely undermine the reliability (and therefore interpretability) of the outcome.

**Figure S1.**
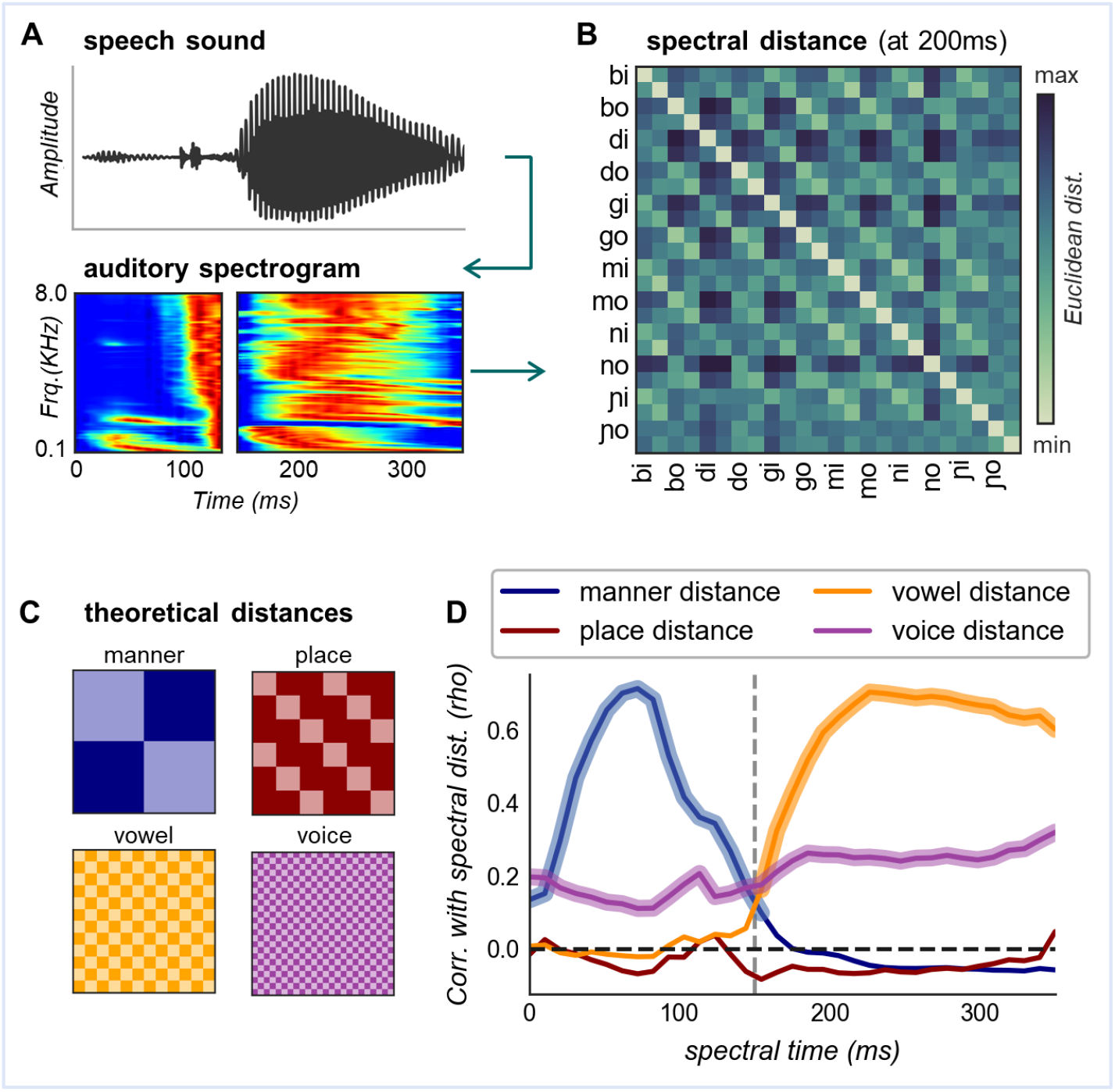
Representational content of the speech stimuli (Figure 1) as they reach the central auditory pathways. (A) Auditory spectrograms were extracted from the speech sounds with a model of cochlear frequency analysis, then averaged by syllable type (top: one instance of “go” pronounced by the female voice; bottom: average spectrogram of all 5 utterances belonging to the sub-condition “go-female”). The blue-red scale reflects minimal-maximal energy, separately normalized in the consonant and vowel portions for mere illustrative purposes. (B) Example of dissimilarity matrix reporting the Euclidean distance between each pair of auditory spectrograms at spectral time=200ms. Each label (e.g. “bi”) indexes two sub-conditions: female and male. (C) Categorical dissimilarity models (conditions are ordered as in the matrix above): light colors indicate correspondence (distance=0) while darker colors signify lack of correspondence (distance=1). (D) Correlation between spectral and theoretical distance matrices as syllable unfolds (the dotted vertical line marks the switch between consonant and vowel). Thicker lines indicate significant time points (p<0.05) after FDR correction. Full methodology description, rationale and complementary results are reported in the supplementary text above.

**Figure S2 (complement of Figure 1).**
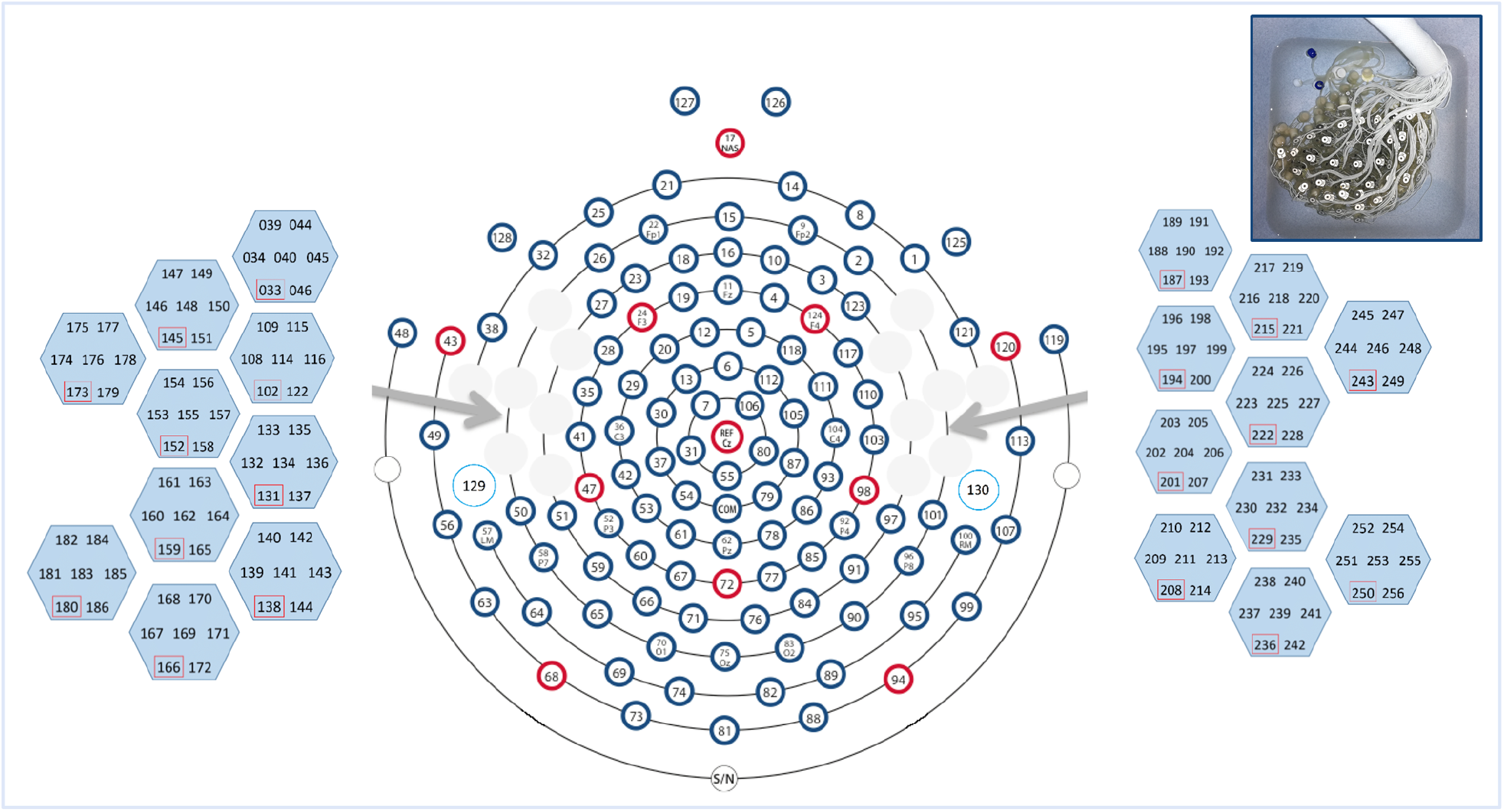
Prototype ultra-high density net. Tight grids of custom electrodes are arranged over the auditory linguistic areas of the superior temporal lobe: 20 temporal geodesic locations (128 partitioning) are filled with hexagonal pods, each containing 7 sensors displaced at a reciprocal distance of 5 mm.

**Figure S3 (complement of Figure 2).**
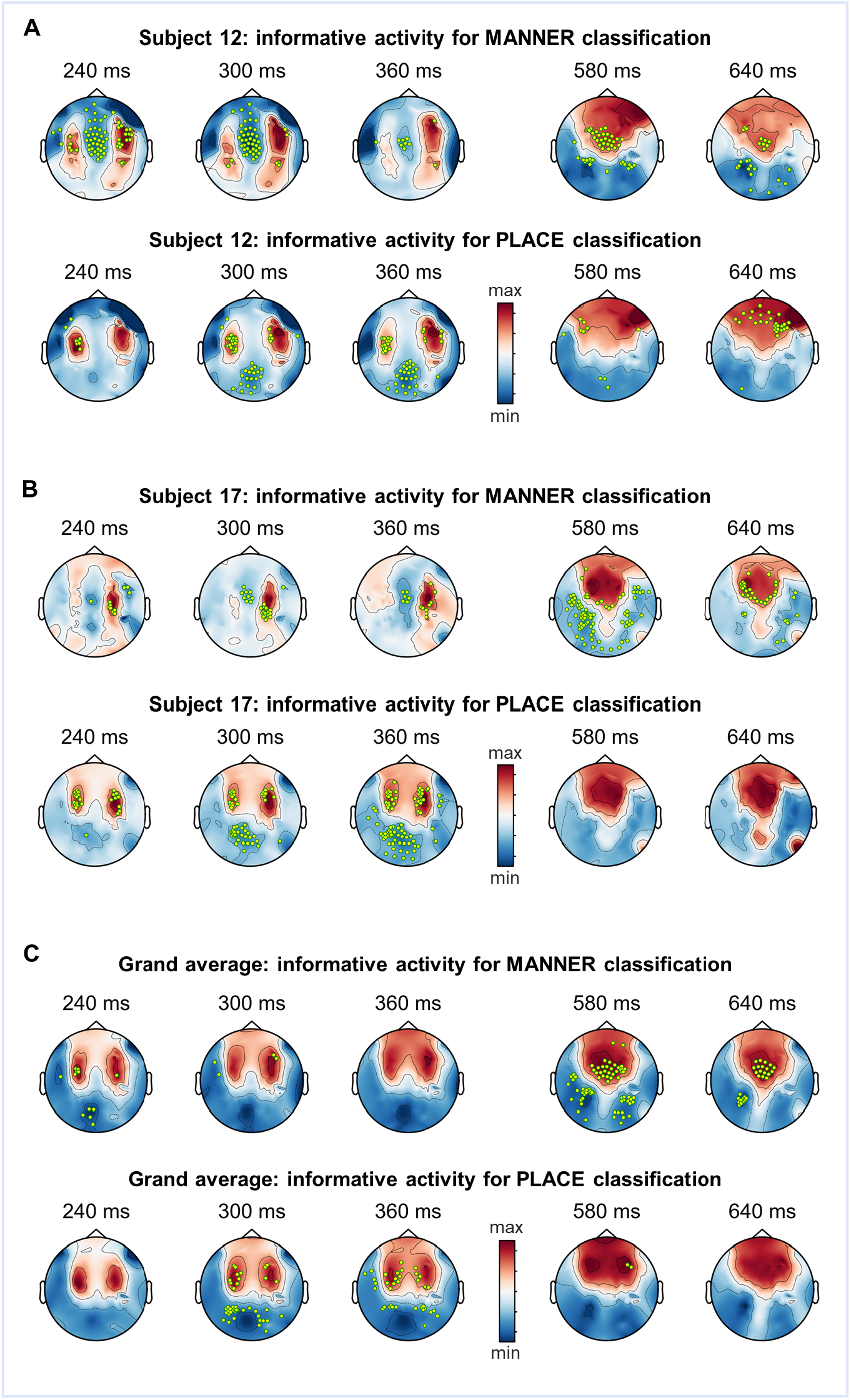
Discriminative loci change as a function of time and phonetic feature dimension. Classifiers weights are projected onto the EEG sensor activation space. Darker colors correspond to brain activity that was useful for classification. Marked in yellow are channels carrying crucial information to distinguish manner but not place (top rows) or to discriminate place but not manner (bottom rows). Time points are chosen to provide an overview of the two time-windows with reliable classification. Panels (A) and (B) show the informative activity patterns reconstructed for two representative subjects. In (C) informative activity patterns are averaged across infants with the purpose of providing a visualization of the general trend. Note however that the interpretability of this grand average is limited since decoding analyses were carried within subject and discriminative loci are very much idiosyncratic. Overall, these topographies show that, as time passes, sensors conveying valuable information are located more medially over frontal areas. Moreover, informative locations for manner and place of articulation do not always overlap.

**Figure S4 (related to Figure 2).**
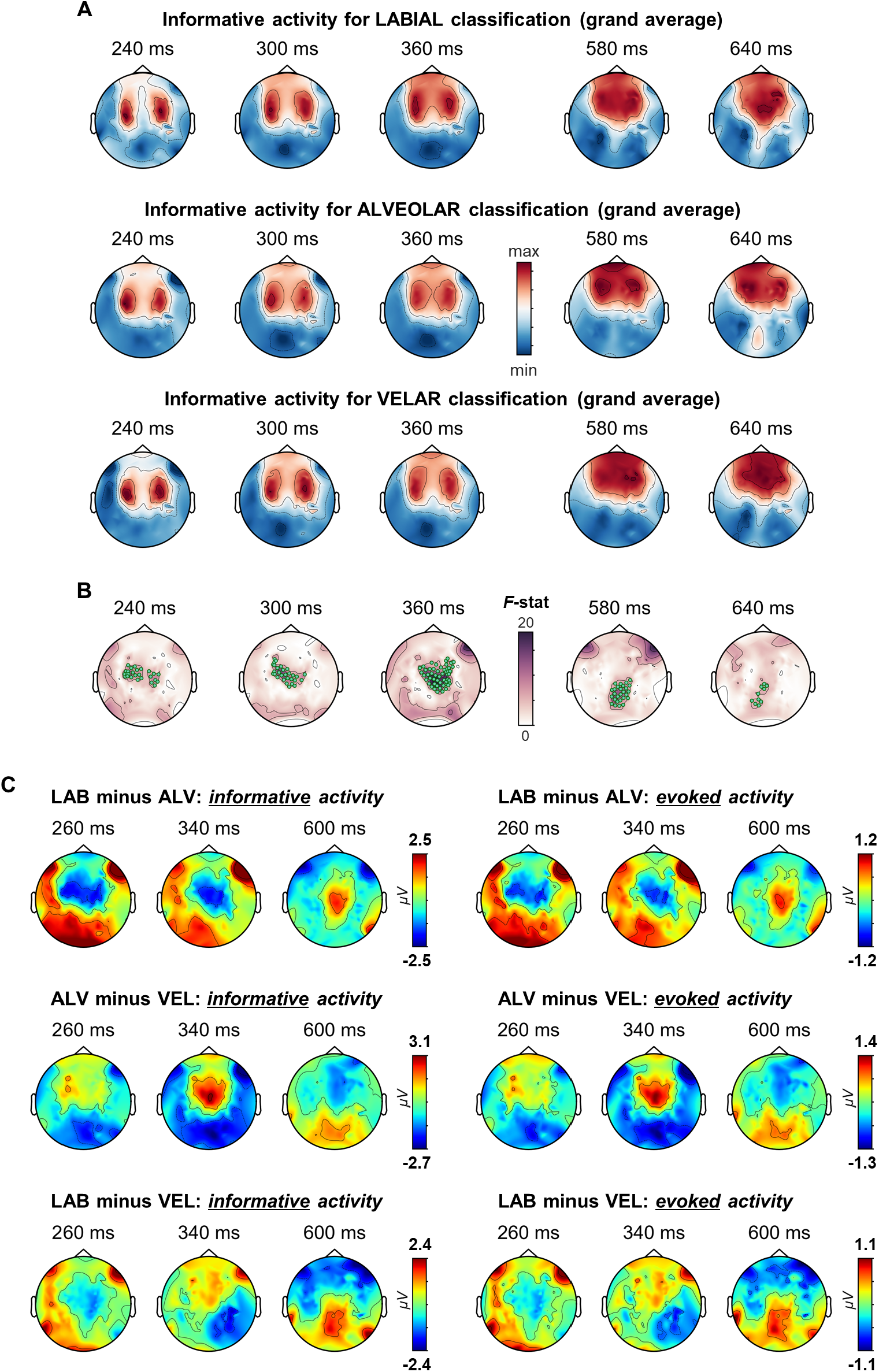
Overview of place contrasts: informative and evoked activity patterns. (A) In place decoding, three distinct models were fitted to separate each place of articulation from the other two (one-vs-rest approach). Their weights were projected back onto the activation space to reconstruct patterns of activity useful in characterizing either labials, alveolars or velars against the other places. Darker colors correspond to loci providing high degrees of class (i.e. place)-specific information. Patterns are averaged across subjects to provide an impression of the general trend; note however that weight idiosyncrasy undermines the interpretability of the grand average. (B) Results of one-way repeated measures ANOVA comparing discriminative activity for labials vs. alveolars vs. velars; channels containing significant differences are in green: early time-window: p_clust_ =0.0005, late time-window: p_clust_ =0.0196. (C) Reported on the left are differential informative activity patterns, on the right the same differences were computed on the evoked related potentials (ERPs). Given that amplitude ranges of informative and evoked brain activity were extremely similar (spanning from −8 to 7 μV in both cases), this figure displays two remarkable features: differential topographies are qualitatively overlapping while amplitude scales (colorbars) change substantially from the left to the right side of the panel.

**Figure S5 (complement of Figure 4).**
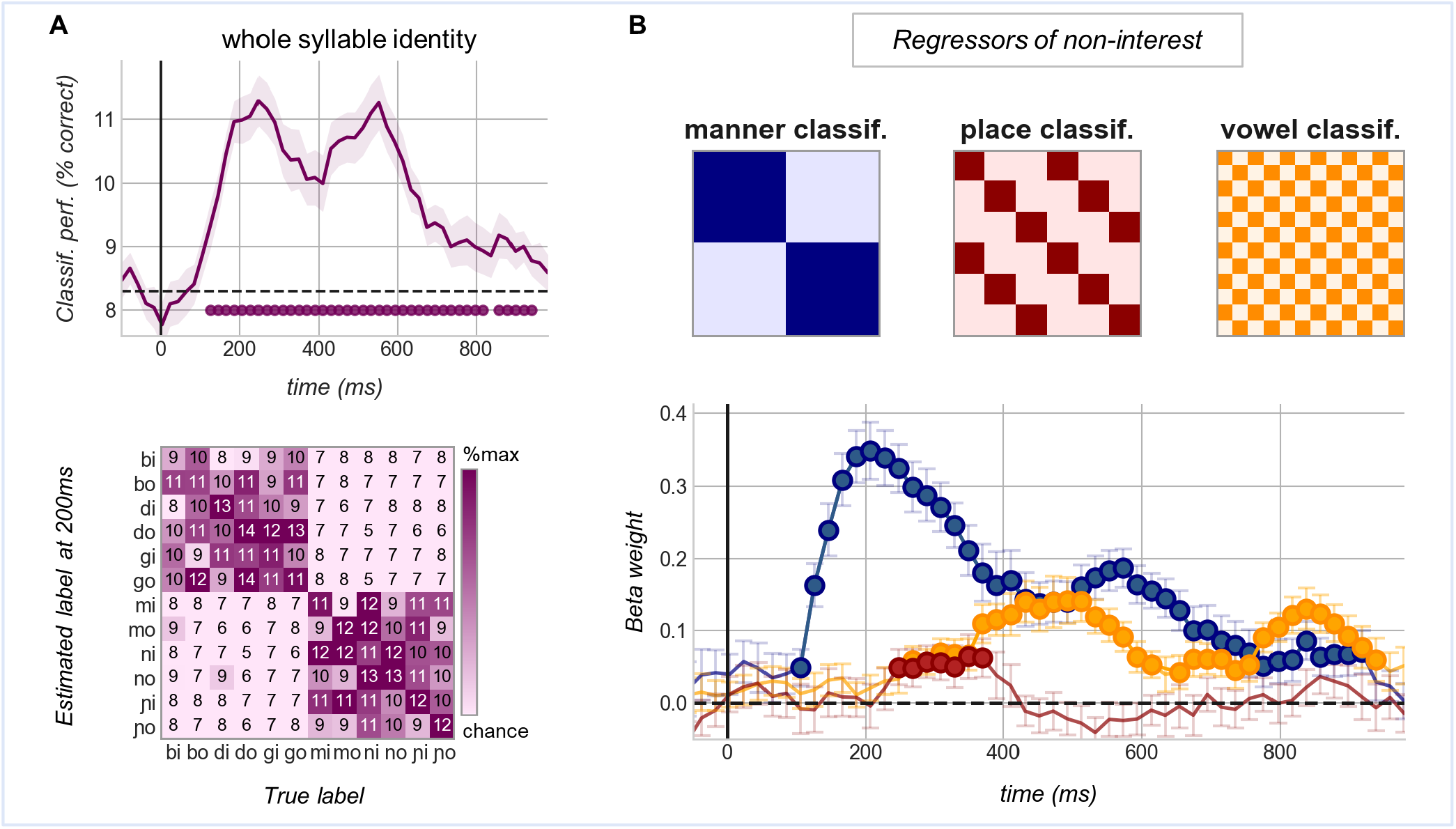
(A) Top: time-resolved accuracy scores of classifiers trained on syllable identities: “bi” vs “bo” vs “di” vs “do” vs “gi” vs “go” vs “mi” vs “mo” vs “ni” vs “no” vs “ɲi” vs “ɲo”. The shaded area corresponds to the SEM across subject, dotted black lines mark theoretical chance level, filled circles indicate when performance is significantly above chance (starting from 120ms: p_clust_=0.0001) Bottom: confusion matrix yielded by the same classifiers at 200ms after stimulus onset. Numbers within each cell indicate the percentage of times a given syllable indicated along the x-axis was classified with the label reported on the y-axis. Off-diagonal values diverging from 0 signal misidentification (chance=8.3%). This example shows how, early on within the trial, classification is mainly driven by manner distinctions. (B) Top: theoretical confusion matrices depicting a perfect separation between (i.e. the ideal classification of) manners of articulation, places of articulation and co-articulated vowel (classes are ordered as in A). Darker colors correspond to the values: 16.6%, 33.3% and 16.6% respectively; light colors correspond to 0%. These matrices were entered as predictors of non-interest in the multiple regression analysis. Bottom: the obtained beta-weights, averaged across subjects and marked by filled circles when significantly above zero (100-920ms: p_clust_=0.0001 for manner; 240-380ms: p_clust_=0.0195 for place, 260-920ms: p_clust_=0.0001 for the vowel). Vertical lines correspond to the SEM.

**Table S1:** Cross-condition decoding. Summary of the decoding performances shown in Figure 3.

| Classes based on: | generalization across: | time window (ms) | p-clust | peak performance |  |  |
| --- | --- | --- | --- | --- | --- | --- |
|  |  |  |  | latency (ms) | score | SD |
| **manner** | speakers | 100-920 | 0.0001 | 200 | 0.673 | 0.079 |
|  | vowels | 100-920 | 0.0001 | 200 | 0.678 | 0.086 |
| **place** | speakers | 200-520 | 0.0001 | 260 | 0.548 | 0.035 |
|  |  | 560-720 | 0.0014 | 640 | 0.522 | 0.047 |
|  | vowels | 240-480 | 0.0001 | 260 | 0.538 | 0.034 |
|  |  | 540-680 | 0.006 | 640 | 0.522 | 0.042 |
| **vowel** | speakers | 260-580 | 0.0001 | 460 | 0.561 | 0.078 |
|  |  | 760-920 | 0.0002 | 800 | 0.554 | 0.052 |
|  | manners | 300-580 | 0.0001 | 460 | 0.57 | 0.08 |
|  |  | 680-960 | 0.0001 | 820 | 0.552 | 0.067 |
|  | places | 280-600 | 0.0001 | 480 | 0.564 | 0.082 |
|  |  | 760-960 | 0.0001 | 820 | 0.544 | 0.066 |

**Table S2.** Formal comparison between main and cross-condition decoding of phonetic features. Performance of estimators trained on exclusive conditions (“across”; Figure 3A-B) is compared to that of estimators trained on all conditions at once (“overall”; Figure 2A-B). AUC scores were averaged over 200ms (the first time point to be considered was set upon peak performance) and, once ascertained the normality of each distribution, contrasted with two-sided t-tests.

| phonetic feature | time window (ms) | decoding analysis | comparison to overall classification |  |  |
| --- | --- | --- | --- | --- | --- |
|  |  |  | mean score | t(24) | p |
| **manner** | 200 - 400 | overall | 0.685±0.065 |  |  |
|  |  | across genders | 0.643±0.057 | 2.278 | 0.032 |
|  |  | across vowels | 0.649±0.0523 | 2.176 | 0.040 |
| **place** | 260-360 | overall | 0.538±0.039 |  |  |
|  |  | across genders | 0.548±0.031 | -1.085 | 0.289 |
|  |  | across vowels | 0.536±0.033 | 0.194 | 0.848 |
|  | 580-680 | overall | 0.526±0.039 |  |  |
|  |  | across genders | 0.513±0.041 | 1.353 | 0.189 |
|  |  | across vowels | 0.519±0.032 | 0.651 | 0.521 |

## Notes

### Competing Interest Statement

The authors have declared no competing interest.

